# Attomol-level quantification of chemically modified ribonucleosides enabled by capillary porous graphitic carbon columns in nano LC-MS

**DOI:** 10.1101/222315

**Authors:** L. Peter Sarin, Sandra D. Kienast, Johannes Leufken, Robert L. Ross, Patrick A. Limbach, Christian Fufezan, Hannes C. A. Drexler, Sebastian A. Leidel

## Abstract

Post-transcriptional chemical modifications of (t)RNA molecules are crucial in fundamental biological processes, such as translation. Despite their biological importance and accumulating evidence linking them to various human diseases, technical challenges have limited the development of methods for reliable detection and accurate quantification of these modifications. Here, we present a sensitive capillary nanoflow liquid chromatography mass spectrometry (nLC-MS) pipeline for quantitative high-resolution analysis of ribonucleoside modifications from complex biological samples. We evaluated two porous graphitic carbon (PGC) materials as stationary phases for reversed-phase separation of ribonucleosides and found that both PGC matrices have excellent retention and separation characteristics, as well as the capability to separate structural isomers. Using PGC matrices in nLC-MS yielded excellent signal-to-noise ratios in a detection range spanning up to six orders of magnitude, allowing for the analysis of individual ribonucleosides down to attomol concentrations. Furthermore, normalizing the obtained signal intensities to a stable isotope labeled spike-in enabled direct comparison of ribonucleoside levels between different samples. In conclusion, capillary PGC columns coupled to nLC-MS constitute a powerful and sensitive tool for quantitative analysis of chemically modified ribonucleosides in complex biological samples. This setup will be invaluable for further unraveling the intriguing and multifaceted biological roles of RNA modifications.

## INTRODUCTION

Translation of the genetic code into functional proteins is fundamental for every living organism. This process is tightly regulated at multiple layers and integrates many players (1) including transfer RNAs (tRNA) – the adapter molecules that link the genetic information to a specific amino acid. These tRNA molecules are decorated by a plethora of evolutionary conserved post-transcriptional chemical modifications (2-4). It has been shown that tRNA modification patterns change in response to physiological and chemical stresses (5,6). These modifications are crucial for fine-tuning translation and thereby maintaining proteome integrity (7-10). Furthermore, a growing number of studies link changes in tRNA modification to various diseases (11,12) including neoplastic proliferation (13), type II diabetes (14), and developmental disorders (15,16). Despite these advances, our understanding of the mechanisms that underlie tRNA modification is still in its infancy. This is in part because quantitative analysis of precious samples, such as patient material of limited availability, requires highly sensitive measurement techniques. Thus, to unravel the biological roles of ribonucleoside modifications, it is absolutely essential to develop techniques that enable their accurate, sensitive, and robust quantification (17-19).

Albeit conceptually straightforward, technical aspects limit the broad application of high-throughput analyses of nucleoside modifications. For instance, high-resolution separation of ribonucleosides by liquid chromatography is challenging due to the very polar nature of the analytes (20). Hence, silica-based C8 and C18 stationary phases often fail to provide satisfying retention and separation characteristics. Densely bonded and end-capped C18 materials are an alternative (6,17,18,21) although they fail to separate closely related compounds, such as positional isomers or nucleoside analogs (22). This limits their applicability for studying complex biological samples and makes a stationary phase with enhanced stereo-selective properties highly desirable.

Porous graphitic carbon (PGC) constitutes an interesting alternative, as it is able to separate highly polar or charged analytes (23,24) and structurally closely related species, such as isomers and enantiomers (25,26). These extended separation and retention characteristics are attributed to its unique structure: PGC consists of planar, two-dimensional sheets composed of hexagonally arranged carbon atoms where each sp^2^ hybridized carbon is joined to three adjacent carbons, resulting in an extensive polyaromatic scaffold. The resulting large delocalized ŏ-electron system yields strong retention of highly polar compounds through ŏ-ŏ electrostatic stacking interactions (27). This polar retention effect, where increasing analyte polarity leads to increased retention, is a characteristic property unique to graphite-based solid phases. Furthermore, PGC utilizes the ability of analytes to adsorb to the planar graphitic surface for separation, providing PGC with a superior stereo-selectivity over phases that mainly rely on hydrophobic interaction between the analyte and the stationary phase (28). Finally, PGC materials are stable over a wide temperature and pH range (27). This has made PGC a popular choice for solid phase extraction (29) and has led to their use in a wide range of analytical applications of biomolecules (30,31), as well as inorganic (32) and organic (33) molecules.

To benefit from these advantages, we sought to develop a robust and reproducible high-resolution nanoflow system to quantify ribonucleoside modifications from complex biological samples. Hence, we tested two PGC stationary phases in a nLC electrospray ionization mass spectrometry (ESI-MS) system to evaluate their suitability for ribonucleoside analysis. We obtained excellent chromatographic characteristics allowing reliable detection of individual ribonucleosides down to attomol concentrations. We showed that PGC materials are able to consistently separate numerous positional isomers, including uridine, pseudouridine, and methylated bases of adenosine, cytidine, and uridine. Finally, we demonstrated that a quantitative analysis of ribonucleoside modification levels is achieved, both label-free and with a stable isotope labeled spike-in, throughout a linear detection range spanning up to six orders of magnitude. To this end, metabolic labeling of ribonucleosides in *Chlamydomonas reinhardtii* constitutes a cheap and facile solution for generating spike-in standards for quantification. Consequently, our methodology is highly suitable for reproducible, quantitative high-resolution analysis of ribonucleoside modifications, and it will significantly contribute to further unravel their biological roles.

## MATERIAL AND METHODS

### Preparation of synthetic ribonucleoside standards

Chemically synthesized mass spectrometry grade ribonucleosides (Carbosynth Ltd, Berkshire, UK; apart from 5-methoxycarbonylmethyluridine (mcm^5^U), 5-methoxycarbonylmethyl-2-thiouridine (mcm^5^s^2^U), 5-carbamoylmethyluridine (ncm^5^U), and 5-carbamoylmethyl-2-thiouridine (ncm^5^s^2^U) that were generously provided by Andrzej Małkiewicz, University of Łódź, Poland) were individually dissolved in 5mM NH_4_HCO_2_ pH 5.3 at a concentration of 50ng/μl. All 4 canonical bases and 28 ribonucleoside modifications (Table 1) were combined in equal amounts to yield the “Complete Standard Mix (CSM)” at a concentration of 1.6μg/μl. A “Limited Standard Mix (LSM)” consisting of 20 bases (Table 1) with a concentration of 1.0μg/μl was also prepared for UPLC analysis.

### Cell culture and metabolic labeling of ribonucleosides

The haploid *Saccharomyces cerevisiae* strain S288C BY4741 was grown in rich growth media (YPD; Formedium) for tRNA isolation. In brief, overnight starter cultures were grown at 30°C, 200rpm. The starter cultures were inoculated into pre-warmed medium to yield an OD_60_0=0.2, and the yeast were grown at 30°C to the logarithmic growth phase (OD_600_=0.8-1.0) and harvested by centrifugation for 3min at 5000*g*.

The cell wall deficient CW15 strain of the green algae *Chlamydomonas reinhardtii* was grown heterotrophically at 22°C with a light intensity of 20μE/m^2^/s and continuous aeration in a rotary shaker at 120rpm for several generations in tris-acetate-phosphate (TAP) medium (34) containing ^15^NH_4_Cl (99.4% ^15^N, Cambridge Isotope Laboratories, Inc., Tewksbury, MA, USA) as the sole nitrogen source. The cells were harvested by centrifugation at 2500g for 5min (35). Non-labeled ribonucleosides were obtained as described above, apart from using NH_4_Cl with a natural isotope distribution.

### tRNA isolation and preparation for LC-MS analysis

Isolation and purification of tRNA from *C. reinhardtii* and *S. cerevisiae* was performed as previously described (6). *C. reinhardtii* tRNA was additionally subjected to gel extraction from a denaturing 8M urea 8% polyacrylamide gel (36). Subsequently, tRNA was enzymatically digested and dephosphorylated to yield monoribonucleosides (6). Briefly, 10μg of tRNA was combined with 40mU of Nuclease P1 from *Penicillium citrinum* (Sigma-Aldrich Biochemie GmbH, Hamburg, Germany) resuspended in 0.2M CH_3_CO_2_Na (sodium acetate) pH 5.3 and 0.1U of Shrimp Alkaline Phosphatase (Thermo Scientific, Schwerte, Germany) in 30μl reactions at 37°C containing 2mg/ml ZnCl_2_ and 1×NEB3 buffer (New England Biolabs GmbH, Frankfurt am Main, Germany) for near neutral reaction conditions, or 20mM CH_3_CO_2_Na pH 5.3 for acidic conditions. After 1.5h the reaction mixture was supplemented with 15μl 0.5M NH_4_HCO_3_ and incubation at 37°C was resumed for 1h. The reaction was terminated by adding 5.0% C_2_HF_3_O_2_ (trifluoroacetic acid; TFA; Sigma-Aldrich Chemie GmbH, Taufkirchen, Germany) in water to a final concentration of 1.0%. The ribonucleosides were purified with HyperSep Hypercarb SPE Spin Tips (Thermo Fisher Scientific GmbH, Dreieich, Germany), dried to completion in a Savant SpeedVac concentrator (Thermo Fisher Scientific GmbH, Schwerte, Germany), and finally resuspended in 5mM NH_4_HCO_2_ pH 5.3.

### Reversed-phase high performance liquid chromatography of ribonucleosides

Nucleoside analysis was performed on two porous graphitic carbon (PGC) matrices; hereafter referred to as PGC-A (Hypercarb 3μm; pore size 250Å; Thermo Fisher Scientific GmbH, Dreieich, Germany) and PGC-B (Prototype PGC 2.1μm; pore size 250Å; MilliporeSigma, Bellefonte, PA, USA). The PGC-A (2.1×150mm) and PGC-B (2.1×100mm) columns were connected to a Knauer PLATINblue UPLC system equipped with an Autosampler 3950, a PLATINBlue T-1 column thermostat, and a MW-1 UV detector (Dr. Ing. Herbert Knauer GmbH, Berlin, Germany). The solvent system consisted of 5mM NH_4_HCO_2_ pH 5.3 (solvent A) and 100% C_2_H_3_N (acetonitrile; ACN; solvent B). The columns were equilibrated in solvent A with 2% B at a flow rate of 0.5ml/min until a stable baseline was achieved. 25μg of the LSM or 2μg of each individual synthetic ribonucleoside was loaded onto the column and washed for 5min using solvent A with 2% B. Separation was obtained by applying a linear gradient from 2% to 98% B at 0.45ml/min for 40min, followed by a wash at 98% B for 5min and regeneration in solvent A with 2% B for 10min, respectively. Conversely, a complex gradient of 2–10% B for 1min, 10% B for 2min, 10–25% B for 25min, 25–50% B for 10min, and 50–98% B for 1min at 0.45ml/min, followed by a wash at 98% B and regeneration in solvent A with 2% B for 5min and 10min, respectively, was also used. The on-column temperature was maintained at 55°C throughout the run, with post-column cooling set to 25°C. Absorption at 254nm was recorded.

The elution order and average retention time for all ribonucleoside modifications was determined from three technical repetitions using automated peak analysis in OpenLAB CDS EZChrom Edition (Dr. Ing. Herbert Knauer GmbH, Berlin, Germany).

### Lifetime performance of porous graphitic carbon in reversed-phase chromatography

The performance of the PGC-A (2.1×150mm) and PGC-B (2.1×100mm) columns were monitored over 30 consecutive runs using 2μg each of pseudouridine (Ψ), 5-methylcytidine (m^5^C), inosine (I), adenosine (A), and 1-methylguanosine (m^1^G). The same chromatographic conditions were applied as described above for the complex gradient. Column performance (e.g. backpressure, signal-to-noise ratio etc.) was monitored from the run parameters and changes in ribonucleoside separation (retention time, peak symmetry and resolution) were determined using automated peak analysis in OpenLAB CDS EZChrom Edition (Dr. Ing. Herbert Knauer GmbH, Berlin, Germany).

### Packing of porous graphitic carbon into fused silica capillaries

PGC-A, and PGC-B were packed into 75μm inner diameter uncoated fused silica capillaries (Z-FSS-075365; Postnova Analytics GmbH, Landsberg am Lech, Germany), to which frits were prepared (37). In essence, 3 parts of potassium silicate 28/30° solution (Kremer Pigmente, Aichstetten, Germany) and 1 part of formamide were mixed thoroughly and the resulting precipitate was spun down at 14,600g for 20s. One end of the silica capillary was immersed for ~10s in the top layer of the polymerization solution, after which the capillary was baked at 100°C for 4h. The newly polymerized frit was conditioned with 100% ACN for 5min.

Next, ~50mg of PGC material was suspended in a 750μl of a 1:1 C_3_H_8_O (isopropanol; IPA):C_2_H_6_O (ethanol) solution and ultrasonicated for 5min in 2ml flat-bottomed glass vial (MilliporeSigma). The vial was tightly connected to the loading bomb and the PGC slurry was injected into the fritted capillary at an initial pressure of ~50bar, followed by a gradual increase to ~100bar. Once the desired bed length was reached (it takes ~8h to reach 550mm), the capillary was connected to a Proxeon EASY nLC-1000 (Thermo Fisher Scientific GmbH, Dreieich, Germany) and the matrix was compacted to its final bed length (500mm) with a flow of 0.1% formic acid at a constant pressure of 500bar.

### Liquid chromatography mass spectrometry of ribonucleosides

LC-MS analysis was performed using a Proxeon EASY nLC Thermo Fisher Scientific GmbH, Dreieich, Germany) online coupled via a self-packed PGC capillary column and an electropolished stainless steel emitter (Proxeon ES542, 30μm×40mm), connected via a MicroTight True ZDV union (IDEX Europe GmbH, Erlangen, Germany), to a Q Exactive mass spectrometer (Thermo Finnigan LLC, San Jose, CA). The column was attached to a Proxeon nano ESI source (Thermo Fisher Scientific GmbH, Dreieich, Germany). Unless otherwise stated, the total load was 100ng for the synthetic ribonucleosides and 250ng for ribonucleosides derived from biological samples. The solvent system consisted of 5mM ammonium formate pH 5.3 (solvent A) and 2:1 IPA:ACN (solvent B). Separation was achieved using a multi-step gradient (0–12% B in 1min; 12–60% B in 60min; 60–100% B in 1min; hold at 100% B for 25min; 100–0% B in 1min; hold at 0% B for 12min) at a flow rate of 250nl/min, with the on-column temperature set to 55°C. The mass spectrometer was run in the positive mode at a resolution of 70.000. Full MS spectra were recorded in profile mode in the scan range from m/z=100-700, with the AGC target value set to 3×10^6^ and the maximum fill time to 50ms. The five most intense ions were selected for fragmentation. MS2 scans were recorded at a resolution of 35.000 in an isolation window of 4.0m/z with the ACG target value set to 2×10^5^ and the maximum fill time to 120ms. Only single charged ions were allowed (M+H^+^). All measurements were carried out in three technical replicates.

### Linear response calibration curves

Dilution of the CSM yield the following concentrations: 1000pg/μl, 500pg/μl, 250pg/μl, 125pg/μl, 62.5pg/μl, 31.25pg/μl, 20pg/μl, 10pg/μl, 5pg/μl, 2.5pg/μl, 1pg/μl, 0.5pg/μl, 0.2pg/μl, 0.1pg/μl, 0.05pg/μl, 0.02pg/μl, and 0.01pg/μl. 4μl of each dilution was subjected to LC-MS analysis in three technical replicates.

### Quantitative analysis of LC-MS data

Manual identification and quantification of ribonucleosides was performed using Qual Browser in the Xcalibur suite (Thermo Fisher Scientific GmbH, Dreieich, Germany). A lookup list containing the chemical formulae of the expected modifications was compiled (ribonucleoside data from MODOMICS) (38), and the corresponding m/z values (Nuc+H^+^) were generated using the isotope-simulation function in Qual Browser. Next, extracted ion chromatograms were created within a tolerance range of ±0.002Da of the theoretical m/z value, and the maximum intensity of the peak (MaxI) was determined using manual peak annotation (Add Peaks). The identity of the ribonucleoside was verified by its retention time and fragmentation pattern in the MS1 and MS2 scans. The MaxI values were normalized against their ^15^N-labeled isoforms in samples spiked with a stable isotope labeled standard.

Semi-automated quantitative data analysis was carried out using pymzML (version 2.0.0) (39) and pyQms (version 0.5.0) (35,40). Prior to quantification, Thermo RAW files were converted to the mzML format (41) using msconvert as part of Proteowizard (version 3.0.10738) (42). Using a manually curated lookup list of all known ribonucleosides (data from MODOMICS) (38), pyQms calculates high accuracy isotopologue patterns derived from their chemical formulas and matches those onto MS1 spectra. Quantification is based on the maximum intensity of the matched isotope pattern chromatogram (MIC) of each ribonucleoside (40). In addition, pyQms generates a weighted similar match score (mScore) to assess the quality of all quantifications (43).

## RESULTS

### Porous graphitic carbon is well suited for separation of polar compounds

PGC materials efficiently retain polar compounds and are capable of separating structurally related substances (44). We sought to test the suitability of PGC materials for reversed-phase separation of chemically modified ribonucleosides using a UPLC system coupled with UV-detection. We then applied this methodology to a nLC system coupled with ESI-MS detection.

The manufacturing of PGC results in a material without precise definition, which prompted us to use PGC materials from two different suppliers. We first established a simple stepwise gradient using narrow bore columns (I.D. 2.1mm) to resolve a mix of 20 chemically synthesized ribonucleosides (LSM, Table 1). Both PGC materials were able to resolve the individual ribonucleosides, with baseline separation obtained for most (Figure 1A; PGC-A and PGC-B). Ribonucleosides having zwitterionic character showed some peak broadening and tailing, attributed to a greater orbital overlap between the delocalized electrons in both analyte and substrate. For most analytes in the sample, peaks were symmetric and signal intensity was high. However, we observed minor variations in resolution and peak symmetry in both PGC materials (Figure 1A). To evaluate these discrepancies between the materials we applied 1μg of each chemically synthesized ribonucleoside onto the PGC columns. Both materials yielded similar elution profiles (Figure 1B) with both having the resolving power to discriminate between positional isomers, such as the methylated adenosines.

**Figure 1.**
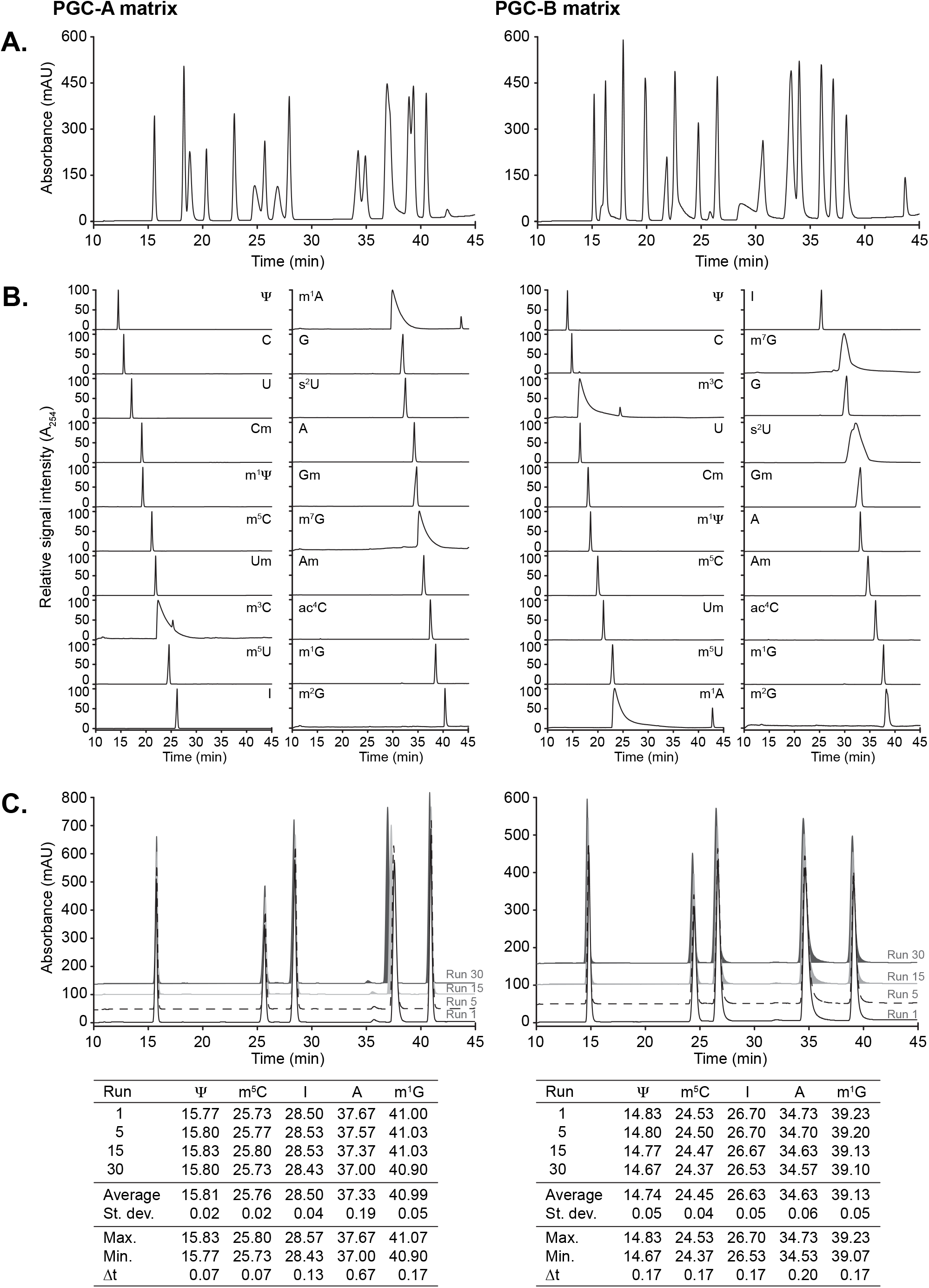
Robust and reliable separation of ribonucleosides is achieved by reversed-phase chromatography using porous graphitic carbon (PGC) materials. (A) Representative chromatograms (A_254_-trace) obtained with PGC-A (left panel) and PGC-B (right panel) separating 20 chemically synthesized ribonucleosides (LSM, Table 1). (B) Individual chromatograms obtained for each of the 20 ribonucleosides included in the LSM using run conditions as in (A). (C) Retention time and column durability analysis of 30 consecutive runs with pseudouridine (Ψ), 5-methylcytidine (m^5^C), inosine (I), adenosine (A), and 1-methylguanosine (m^1^G) on PGC-A and PGC-B columns. Representative chromatograms from runs #1, #5, #15, and #30 are shown. The table summarizes the retention time parameters for all tested nucleosides. Abbreviations follow the Modomics database convention (38).

Nonetheless, we did observe some interesting differences. First, certain ribonucleosides change their elution order, as 3-methylcytidine (m^3^C) elutes as nucleoside number eight using PGC-A, but as nucleoside number three using PGC-B. More subtle differences are seen for 2’-O-methylguanosine (Gm), 7-methylguanosine (m^7^G), and 2-thiouridine (s^2^U) that change by one or two positions between the materials (Figure 1B, Table 2). Second, 1-methyladenosine (m^1^A), m^3^C, and m^7^G yield broad peaks in both PGC materials (Figure 1B, Table 2), although PGC-B is more strongly affected. Third, the signal intensities differ somewhat between PGC-A and PGC-B. This is particularly apparent for s^2^U, but also m^7^G and, to a lesser extent, other methylated guanosines (Table 2). Fourth, PGC-B separates most uridine analogs very well, with the exception of thiolated moieties, yielding highly symmetric peaks with strong intensity (Table 2). Hence, PGC materials offer a powerful strategy for robust chromatographic resolution of modified ribonucleosides, albeit it is critical to evaluate material-specific nuances for targeted applications.

### PGC materials are robust for long-term applications

PGC materials are chemically inert, withstanding acidic or basic conditions and high temperatures without undergoing degradation as seen in silica based stationary phases. However, hyper-retention can cause a buildup of analytes leading to a loss of resolution and increased backpressure (28,45,46). To test the longevity of the PGC materials, we subjected the columns to successive elution and regeneration cycles whereby a mix of five ribonucleosides (Ψ, m^5^C, I, A and m^1^G) was loaded. These ribonucleosides elute at different stages of the gradient, allowing us to monitor hyper-retention by analyzing changes in elution time, peak shape, and resolution. Following a series of 30 consecutive runs, we observed only negligible shifts in retention time for all five analytes (Figure 1C). The largest shift is observed for adenosine in PGC-A, with a difference between the earliest and latest elution time (Σt) of only 0.67min (Figure 1C). Since the theoretical plate counts remain stable (Table 2), this establishes that the chromatographic performance of the PGC matrices is unaltered throughout the assay.

### Self-packed porous graphitic carbon capillary columns provide strong advantages for nanoflow ESI-MS

Sample limitation is often a challenge for quantitative analysis of modified ribonucleosides in biological samples. Downscaling the chromatographic setup to capillary columns coupled with ESI-MS can circumvent this limitation by significantly reducing the amount of input material required. However, the microfluidic environment of capillary columns differs from traditional sized columns, affecting the overall performance of the matrix and preventing direct downscaling. To ensure conformity and comparability, we prepared columns by tightly packing the two PGC materials into silica capillaries (I.D. 75μm), trimmed to a final length of 500mm. When loading 100ng of the LSM (Table 1) we achieved an excellent signal-to-noise ratio even though peaks were less distinct than in the UPLC setup (signal intensities 3.06×10^10^ and 4.07×10^10^; Figure 2A). To evaluate the chromatographic performance of the PGC nano columns in more detail, we generated extracted ion chromatograms (XIC) for the canonical bases and Ψ, I, N4-acetylcytidine (ac^4^C), m^1^A, and 2’-O-methyladenosine (Am) (Figure 2B), and confirmed the identity of the peaks by MS1 and MS2 spectrum analysis (Figure 2C). Both PGC materials displayed high peak intensity and acceptable peak symmetry apart from guanosine, which eluted over a broad range (Figure 2B). Despite the limitations observed for guanosine and its analogs, the PGC materials performed excellent at separating most ribonucleosides (Supplementary Table 1). In a direct comparison, we observe a slightly better resolution and peak symmetry for the PGC-B material, which can be attributed to the smaller particle size and thereby more efficient packaging of the material. Therefore, we proceeded with the PGC-B material and tested its performance on complex biological samples. We applied 100ng of either the defined CSM (Table 1) or an enzymatic digest of total yeast tRNA to the PGC-B column and monitored the elution profiles (Figure 3). As expected, the CSM yields a less complex TIC with readily distinguishable peaks while the yeast tRNA digest leads to a less defined pattern where only a handful of peaks can be distinguished (Figure 3A). Thus, we generated XICs of all detected ribonucleosides from both analytes (Figure 3B,C). Regardless of the complexity of the samples, the column separated most ribonucleosides at high resolution despite strong similarities in their structure and chemical properties (Figure 3B), as well as significant abundance differences in the tRNA sample. We found that ribonucleosides with overlapping retention times in our setup all have different molecular masses, which allows for their accurate identification and quantification. More importantly, ribonucleosides with identical molecular masses can be readily assigned. For example, the methylated cytidines m^3^C, Cm, and m^5^C, which are often elusive on traditional C18 materials, are baseline separated (Figure 3C). Furthermore, the PGC material is capable of separating between acetylated, formylated, and methylated forms of the same base, as demonstrated by f^5^Cm and ac^4^C in the yeast tRNA digest (Figure 3C). Separation is also achieved for methylated adenosines (m^1^A, Am, m^6^A), as well as for uridine, Ψ, and their methylated analogs, but not for the methylated guanosines (m^1^G, m^2^G, Gm, m^7^G) that fail to elute as individual peaks (Figure 3C). This hyper-retention of the methylated guanosines is attributed to the high electronegative density of the molecules as their delocalized electrons interact with delocalized electrons at the PGC surface, forming strongly binding non-covalent interactions between the analyte and the substrate (47,48).

**Figure 2.**
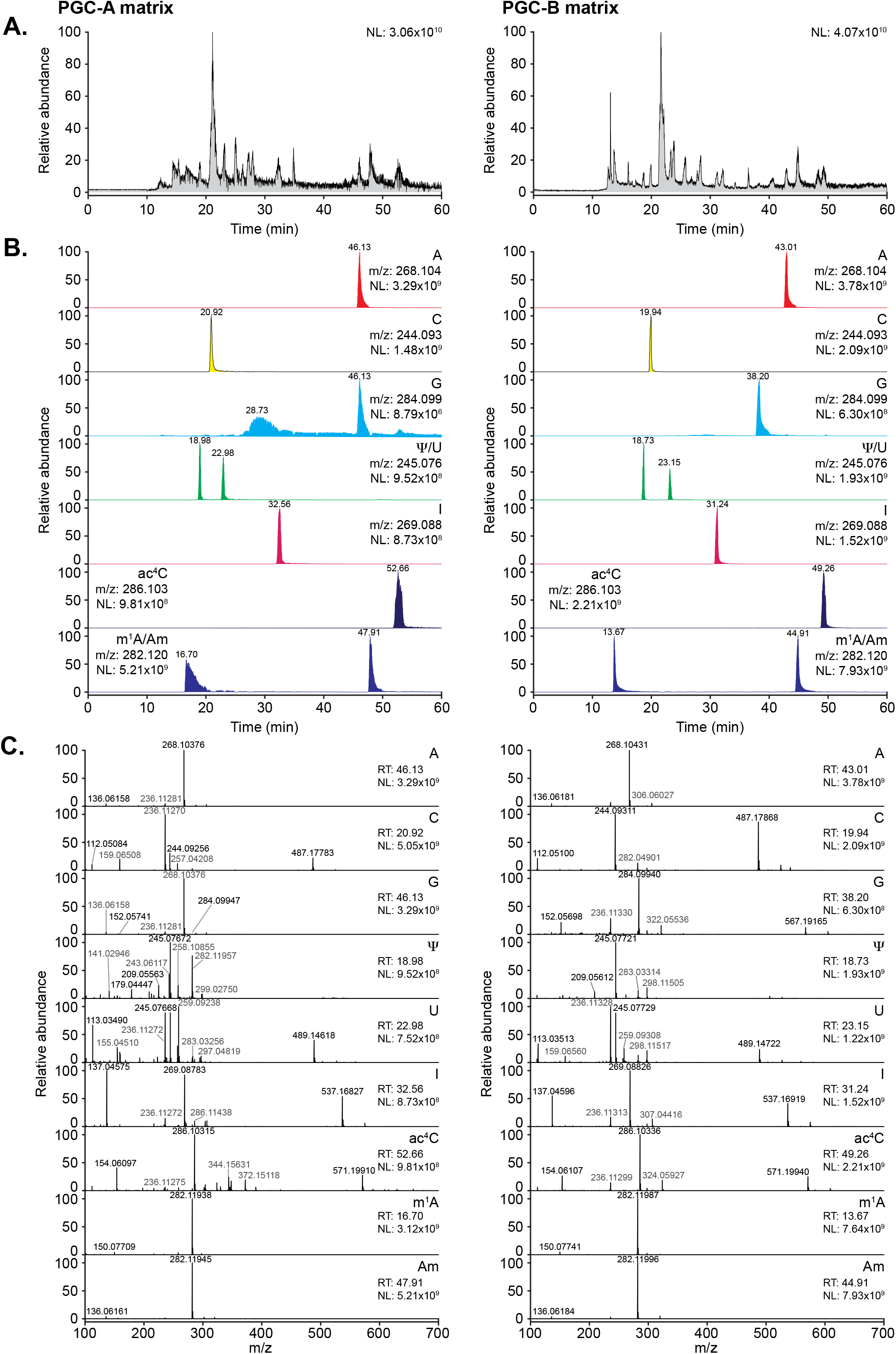
Porous graphitic carbon enables the analysis of ribonucleosides in nLC ESI-MS/MS. A) Representative total ion chromatograms (TIC) of 100 ng of the LSM analyzed on nanoflow capillary columns packed with PGC-A (left panel) or PGC-B (right panel) material. (B) Extracted ion chromatograms (XIC) for adenosine, cytidine, guanosine, uridine, pseudouridine (Ψ), inosine (I), N4-acetylcytidine (ac^4^C), 1-methyladenosine (m^1^A), and 2’-O-methyladenosine (Am). (C) Example MS1 spectra recorded at the retention times corresponding to the nucleosides analyzed in (B). NL: Normalized target level.

**Figure 3.**
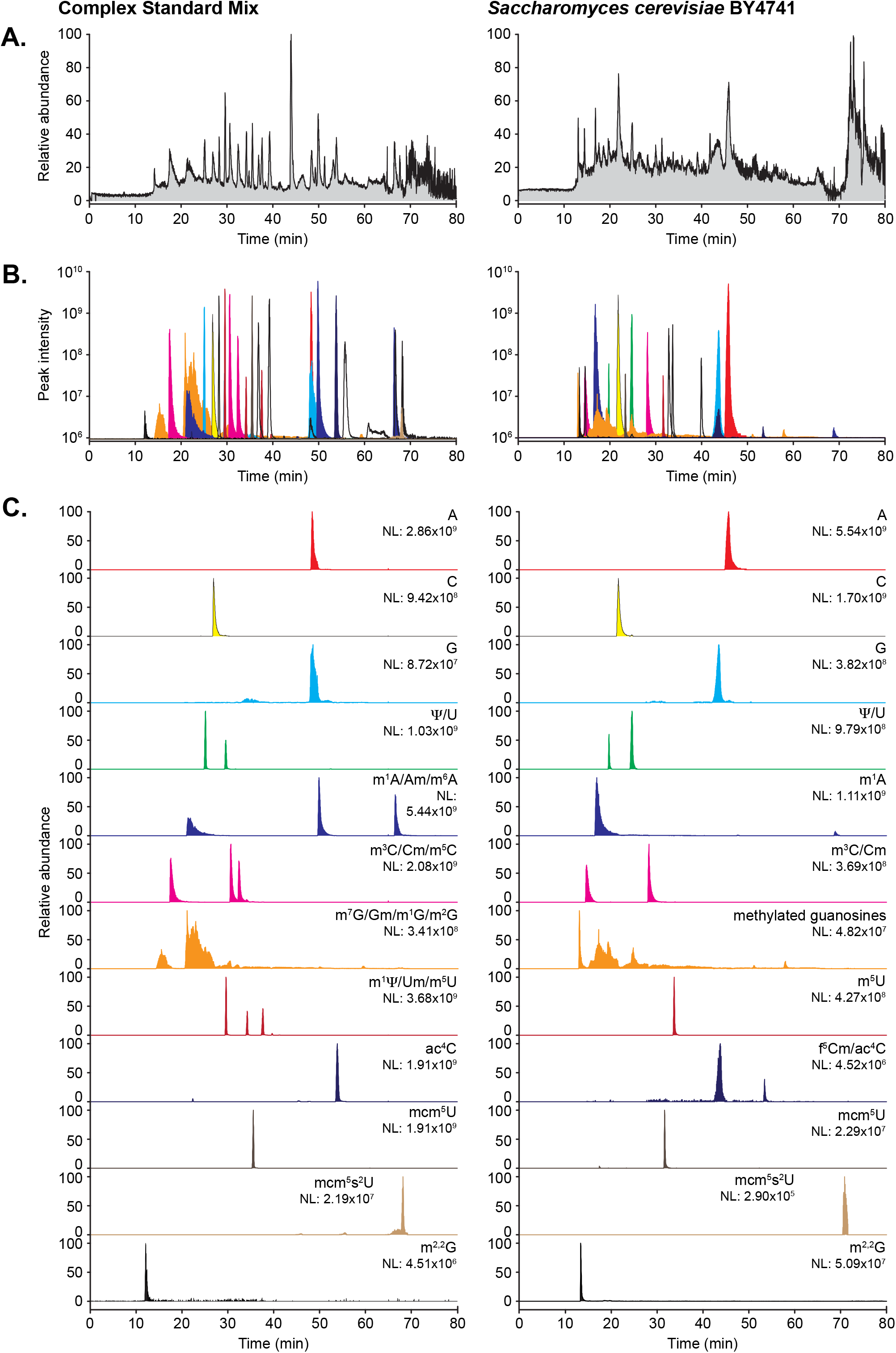
Comprehensive analysis of highly complex analytes is achieved using PGC-B. (A) Representative TIC chromatograms of 32 chemically synthesized ribonucleosides (100ng of CSM, Table 1; left panels) and 250ng of an enzymatic digest of bulk tRNA isolated from *Saccharomyces cerevisiae* strain BY4741 (right panels) analyzed on a nano PGC-B capillary column. (B) Overlay of the XICs for all CSM ribonucleosides detected in the respective samples. (C) XICs of selected nucleosides present in the samples. Note the capability of the PGC material to separate between positional isomers with identical molecular masses. NL: Normalized target level.

Although methylated guanosines are poorly distinguishable from each other, employing PCG materials in nLC ESI-MS provides excellent resolution and high sensitivity for all other ribonucleosides tested. Indeed, weak signals in the range of 10^4^-10^5^ can still be detected, as shown by e.g. mcm^5^s^2^U (Figure 3C), which implies a dynamic range of at least five orders of magnitude for the accurate detection of ribonucleosides.

### Nanoflow porous graphitic carbon capillary columns enable characterization of ribonucleosides at attomol levels

To assess the detection limit and linear range of quantification in our nLC ESI-MS setup, we systematically quantified the maximum peak intensity obtained for each of the 32 ribonucleosides present in the CSM with sample loads spanning six orders of magnitude from 40fg to 4ng (Figure 4). Next, we determined the linear correlation of the maximum peak intensity as a function of the amount of sample loaded. We were able to quantify most of the canonical and modified ribonucleosides within a linear range spanning several orders of magnitude (Figure 4A,B). As expected, guanosine exhibits only a narrow window of less than three orders of magnitude in which it can be reliably quantified (Figure 4A,B). Furthermore, thiolated moieties ionize poorly (Figure 3C) and are moderately resolved by the PGC material (Figure 1C). This results in a poor detection limit of 0.1-0.5ng (~0.3-1.6pmol) and a narrow quantification window (Figure 4B) comparable to C18 materials (6). In contrast, most other ribonucleosides have detection limits lower than 40fg (~100amol), although linear correlation is usually achieved at slightly higher loads of 0.2-10pg (~0.6-30fmol) or above (Figure 4B). In optimal conditions, ribonucleosides can be detected and characterized over a very broad range exceeding six orders of magnitude, whereas the linear range that allows for reliable quantification is significantly smaller.

**Figure 4.**
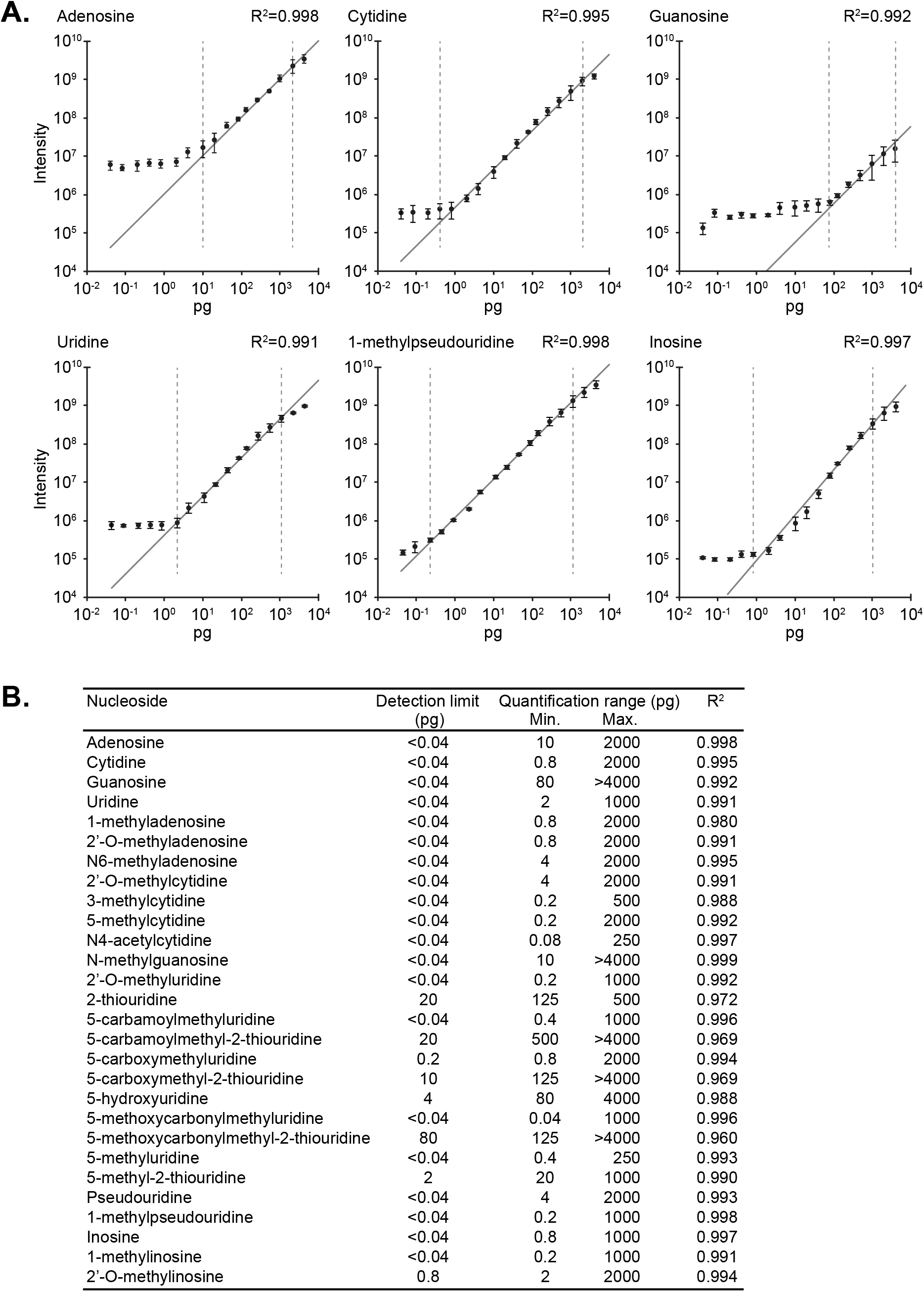
Absolute quantification can be accomplished over a broad detection range. (A) Calibration curves for representative ribonucleosides showing the observed XIC maximum intensity as a function of sample loaded (0.04–4000pg). The error bars represent the standard deviation for each data point (n=3). Linear regression (dark gray line) is used to determine the dynamic range of the instrument for each ribonucleoside by observing the range at which a linear dependency between input amount and intensity is observed (vertical dashed bars indicate the range). (B) Summary of the linear quantification range determined for all 32 ribonucleosides present in the CSM.

### Metabolic labeling enhances quantitative analysis of ribonucleosides

Ribonucleosides can be readily detected and characterized from various biological samples using our nLC ESI-MS setup. Nonetheless, accurate quantitative analysis requires the presence of an invariable internal standard, such as an artificial nucleoside analog (49), to which the modification levels are normalized. However, differences in structure and charge of the ribonucleosides significantly affect ionization, which might bias results if quantifications are normalized only to a single nucleoside. To circumvent this, stable isotope (^15^N)-labeled ribonucleosides can be used as internal standards (Figure 5) (50,51). This allows each modification in the sample to be specifically normalized against its heavier ^15^N-counterpart, thereby eliminating any variance arising from differences in ionization efficiency (Figure 5A-C).

**Figure 5.**
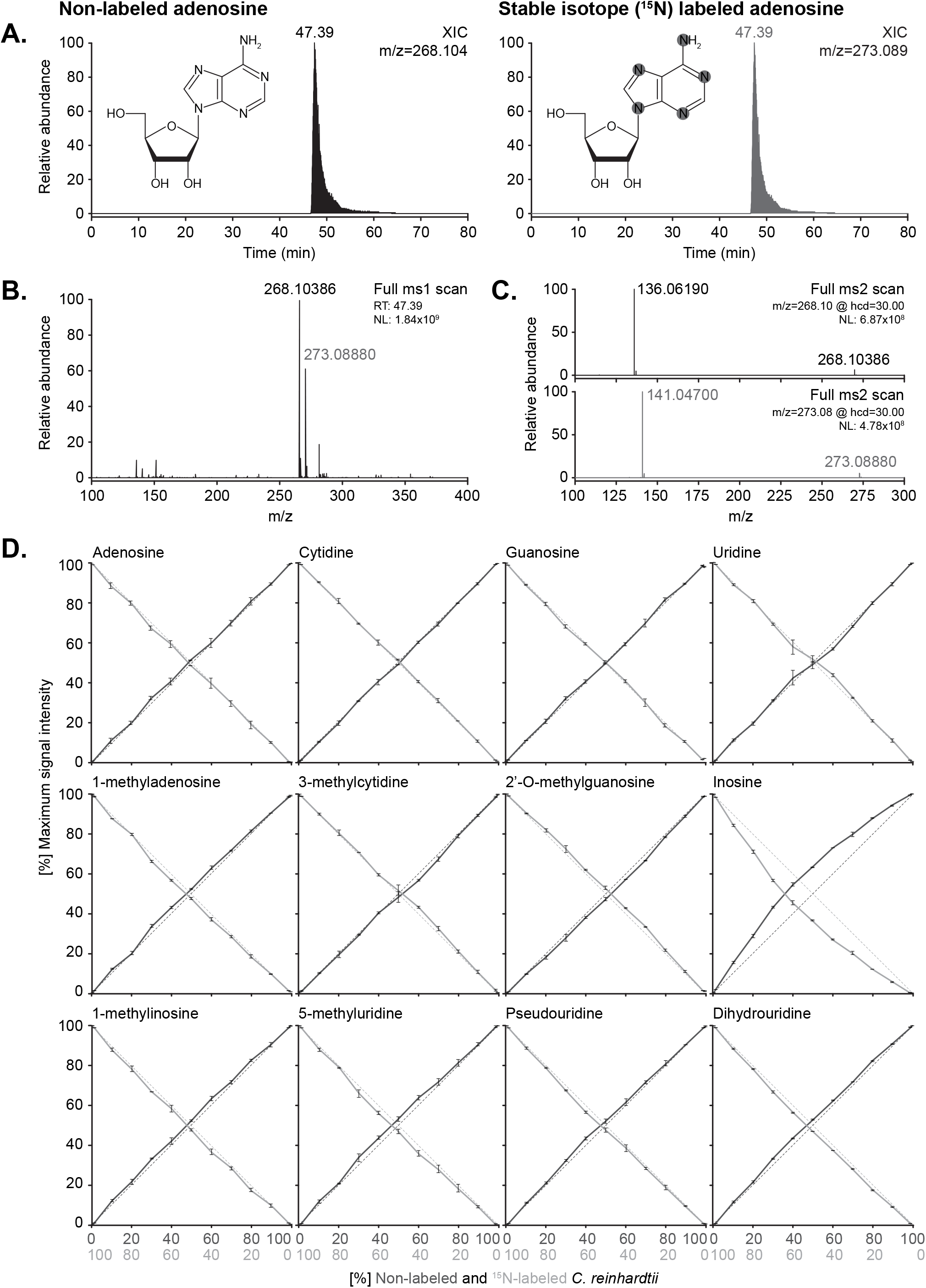
Accurate relative quantification of ribonucleosides can be achieved using stable isotope labeled internal spike-in standards. (A) XICs of adenosine in its native non-labeled (left panel) form and with stable isotope (^15^N)-labeling (right panel, spike-in standard), as well as a schematic representation of the chemical structure of adenosine showing the ^15^N incorporation sites highlighted in gray. (B) MS1 spectrum for adenosine at RT 39.30 min. Note the increase in mass (~5 Da) resulting from the incorporation of ^15^N in the base (^15^N-labeled mass in gray, non-labeled mass in black). (C) MS2 fragmentation spectra of non-labeled (left panel; m/z=268.10) and ^15^N-labeled (right panel; m/z=273.09) adenosine. Note the appearance of the base at m/z=136.06 and m/z=141.05, respectively, corresponding to the expected neutral loss of ribose (-132.04). (D) Cross-dilution series of non-labeled and ^15^N-labeled enzymatic digests of bulk tRNA isolated from *Chlamydomonas reinhardtii*. Quantification of the maximum peak intensity of non-labeled (solid dark gray line) vs. ^15^N-labeled (solid light gray line) XICs of the canonical bases and representative ribonucleoside modifications. Shown is the abundance ratio of non-labeled (N) and ^15^N-labeled (^15^N) maximum signal intensities (MaxI); Abundance = MaxI_15N_/(MaxI_15N_ + MaxI_N_) or vice versa. The dotted lines represent the expected ratio of ^15^N-labeled (light gray) vs. non-labeled (dark gray) material present in the samples. NL: Normalized target level.

Since metabolic labeling of yeast requires a set of expensive reagents, we devised a cheap and simple strategy to generate metabolically labeled spike-in standards. To this end, we grew the single-celled green algae *C. reinhardtii* in a ^15^N-containing growth media (35), achieving full labeling within a few days. To test the suitability of stable isotope labeled spike-in standards for normalization and to simulate a dataset that corresponds to SILAC labeling of ribonucleosides, we devised a cross-dilution series where non-labeled and ^15^N-labeled ribonucleosides isolated from *C. reinhardtii* are combined in specific ratios ranging from 100:0 to 0:100 (Figure 5D). The ^15^N-labeled ribonucleosides yielded a strong signal throughout the cross-dilution series. As little as 2.5ng (99:1 ratio) of spike in standards were sufficient to achieve reliable detection and quantification when mixing with 99-fold more concentrated non-labeled ribonucleosides (247.5ng load). Moreover, the intensities observed throughout the cross-dilution series (Figure 5) were within the linear range of the instrument (Figure 4).

Hence, we analyzed a total of 19 ribonucleosides semi-automatically using pyQms (40) and manually using QualBrowser, yielding identical outcomes (Supplementary Figure 1, Supplementary Table 2). The ratio of non-labeled and labeled ribonucleosides was calculated to determine the relative proportion of each isoform in the cross-dilution series. Importantly, the measured relative abundance of the isoforms accurately reflects the theoretical cross-dilution ratio for almost all ribonucleosides (Figure 5D, Supplementary Figure 1; Supplementary Table 2). Interestingly, for m^5^C, Cm, and I the point of equal intensity between the non-labeled and ^15^N-labeled isoform shifted significantly towards a higher ratio of the non-labeled isoform (Figure 5D, Supplementary Figure 1). As the reproducibility of the measurements (n=3) following normalization to the ^15^N-labeled isoform is very high (m^5^C ±0.62%; Cm ±0.83%; I ±0.42%), it is highly unlikely that technical issues caused this difference. This suggests that the observed shift stems from marginally different amounts of m^5^C, Cm and I in the two samples. Since the *C. reinhardtii* cultures yielding the metabolically ^15^N-labeled and the non-labeled tRNA samples were generated six months apart of each other, they may have encountered slightly different growth conditions that resulted in the observed changes to the modification pattern.

Hence, this shows that SILAC-type experiments are feasible for ribonucleoside analysis. Furthermore, we are able to reliably detect minute changes in modification levels from complex biological samples. This highlights the need of spike-in standards as a powerful tool for normalization.

## DISCUSSION

PGC materials combine several attractive characteristics that make them well suited as column matrices for reversed-phase chromatography of chemically modified ribonucleosides. In particular, the polar-retention effect and the strong stereoselectivity due to planar surface adsorption are unique features that allow for robust separation of complex biological samples, differentiating PGC materials from conventional C18-based stationary phases (23).

In this study, we demonstrate that chemically modified ribonucleosides can be reliably baseline separated at high peak capacity in microbore (Figure 1) and capillary nanoflow (Figure 2) column format using different PGC stationary phases. It is straightforward to establish separation as even a simple linear gradient and different buffer systems yield satisfying results (Supplementary Figure 2). Although the two PGC materials are very similar in their chemical and physical properties, slight differences in retention time, resolution, peak shape, and peak symmetry are apparent (Figure 1B, 2B). This is most likely attributable to differences in physical parameters, e.g. different particle sizes for the PGC matrices (3μm vs. 2.1μm) and different column lengths in the microbore setup (150mm vs. 100mm). Importantly, we did not observe a buildup of analytes due to hyper-retention, which would lead to a gradual decrease in chromatographic performance (27,46). Both PGC matrices remained stable throughout thirty consecutive runs, showing only minute changes in retention time and resolution, as well as only slight fluctuations in peak intensity (Figure 1C). This shows that both PGC materials maintain uniform long-term chromatographic performance.

In the nLC setup, the smaller particle size of PGC-B (2.1μm) shows greater peak capacity compared to PGC-A (3.0μm) albeit at the cost of increased backpressure. However, the strong binding characteristic of both PGC materials allows challenging uridine analogs to be resolved at a high resolution. This makes PGC matrices an attractive choice for the analysis of complex biological samples, since wobble position uridines (U_34_) are frequently modified and known to be critical in different biological scenarios. Indeed, we successfully baseline separated more than 25 ribonucleosides from both synthetic and complex biological samples on PGC capillary columns, achieving excellent signal-to-noise ratios in a detection range that spans up to six orders of magnitude (Figure 3,4). However, the four methylated guanosines (m/z=298.118) included in the CSM (m^1^G, m^2^G, m^7^G, Gm) remain poorly resolved on both PGC matrices (Figure 3C), although guanosine itself has a slightly limited resolution (Figure 2B, 3C). Conversely, N2,N2-dimethylguanosine (m^2,2^G), a double methylated form of guanosine, was detected without difficulty at high signal intensity in the biological sample (Figure 3C). A striking key advantage of PGC columns is their ability to separate ribonucleosides with identical molecular weights (e.g. m^3^C, m^5^C, and Cm) using simple mobile phase gradients. This enables reliable quantification without the need for MS2 or MS3 fragmentation strategies. Nevertheless, it is crucial to use ribonucleoside standards to determine the elution order as it may shift slightly between different PGC materials.

In complex biological samples, detecting the ribonucleoside of interest is merely the first step, as crucial information about its function and regulation can only be deduced from its abundance. This necessitates a quantitative approach whereby absolute or relative abundances of the analytes can be precisely determined by the use of calibration curves, spike-in standards, and a fine-tuned instrument with a broad dynamic range. By determining the limit of detection and linear range for each ribonucleoside, it is possible to set up the mass spectrometer to measure absolute abundances. Indeed, we found that most of the ribonucleosides studied have a linear range spanning three to four orders of magnitude (Figure 4). Within this range, we can determine absolute quantities of ribonucleosides present in any given sample by correlating maximum peak intensity to sample amounts. The major caveat in this approach is the need for synthetic nucleosides, i.e. each ribonucleoside of interest can only be quantified once the detection limits of the instrument have been determined using a pure synthetic homolog (Figure 4). Even though the number of synthetic nucleosides that are commercially available is increasing, absolute quantification of many modifications requires expensive custom-synthesized standards. However, for most biological applications the need for synthetic nucleosides can be circumvented, as the change in relative abundance between samples is more informative than absolute quantification. Here, stable isotope (^15^N-)labeled ribonucleoside standards from *C. reinhardtii* provide an invariable internal reference point to which different samples can be compared and normalized (Figure 5, Supplementary Figure 1). Using *C. reinhardtii* provides a simple source of ^15^N-labeled ribonucleosides that are approximately 10–15 times cheaper than those isolated from ^15^N-labeled yeast. This circumvents metabolic labeling of biological samples in cases where it is not always straightforward, e.g. due to the high costs associated with defined isotope-labeled growth media, or in the cases where it is simply not possible, such as clinical samples. As we have successfully demonstrated, even low amounts of metabolically ^15^N-labeled spike-in standards yield sufficiently strong signals for reliable normalization and quantification (Figure 5D). Moreover, the technical reproducibility of measurements from the same biological sample can be significantly improved by internal normalization. For example, the maximum peak intensity recorded for A, C, G, and U in three technical repeats yields 10.6%, 10.0%, 12.2%, and 9.4% standard deviation prior to normalization, but merely 0.89%, 0.46%, 0.57%, and 0.78% following normalization to the ^15^N-isoform. This constitutes a 12 to 22-fold increase in precision. Furthermore, subtle changes in culture conditions may lead to changes in individual tRNA modifications, which can be detected by the use of labeled ribonucleosides in SILAC-like experiments. Finally, we established that pyQms (40) is a robust tool for semi-automated MS based ribonucleoside quantification, achieving identical results to manual quantification in a fraction of the time required for analysis.

In conclusion, we have shown that a nLC ESI-MS setup coupled to capillary PGC nano columns can be successfully used to quantitatively analyze chemically modified ribonucleosides. The excellent resolution and strong signal-to-noise ratio obtained for, in particular, pyrimidine-based ribonucleosides makes PGC an excellent chromatographic matrix for the study of complex biological samples. Furthermore, combining this chromatographic setup to a MS detector with improved sensitivity could further lower the detection boundary towards zeptomol amounts and expand the linear range for quantification.

## AVAILABILITY

Modomics is a database of RNA modification pathways available at http://modomics.genesilico.pl. pymzML (version 2.0.0) is an open source python module for high-throughput bioinformatics analysis of mass spectrometry data, and pyQms (version 0.5.0) is an open source python module for universal and accurate quantification. Both modules are available in the GitHub repository (https://github.com/pymzml/pymzML; https://github.com/pyQms/pyqms). Proteowizard (version 3.0.10738) is an open source, cross-platform tool and library collection for proteomics data analyses available in the Sourceforge repository (http://proteowizard.sourceforge.net).

## ACKNOWLEDGEMENT

The authors wish to thank Dr. Daniel Shollenberger and Dr. David Bell at MilliporeSigma for kindly providing PGC material, Prof. Michael Hippler for *Chlamydomonas* material and media, Annalen Nolte and Karin Scharmann for technical support, and members of the Leidel lab and SPP1784 for critical discussions. R.L.R. thanks the University of Cincinnati RITE program for supporting his international travel and research experience.

## TABLE AND FIGURES LEGENDS

**Table 1**. Synthetic ribonucleosides used in this study.

**Table 2**. Chromatographic properties of microbore PGC columns.

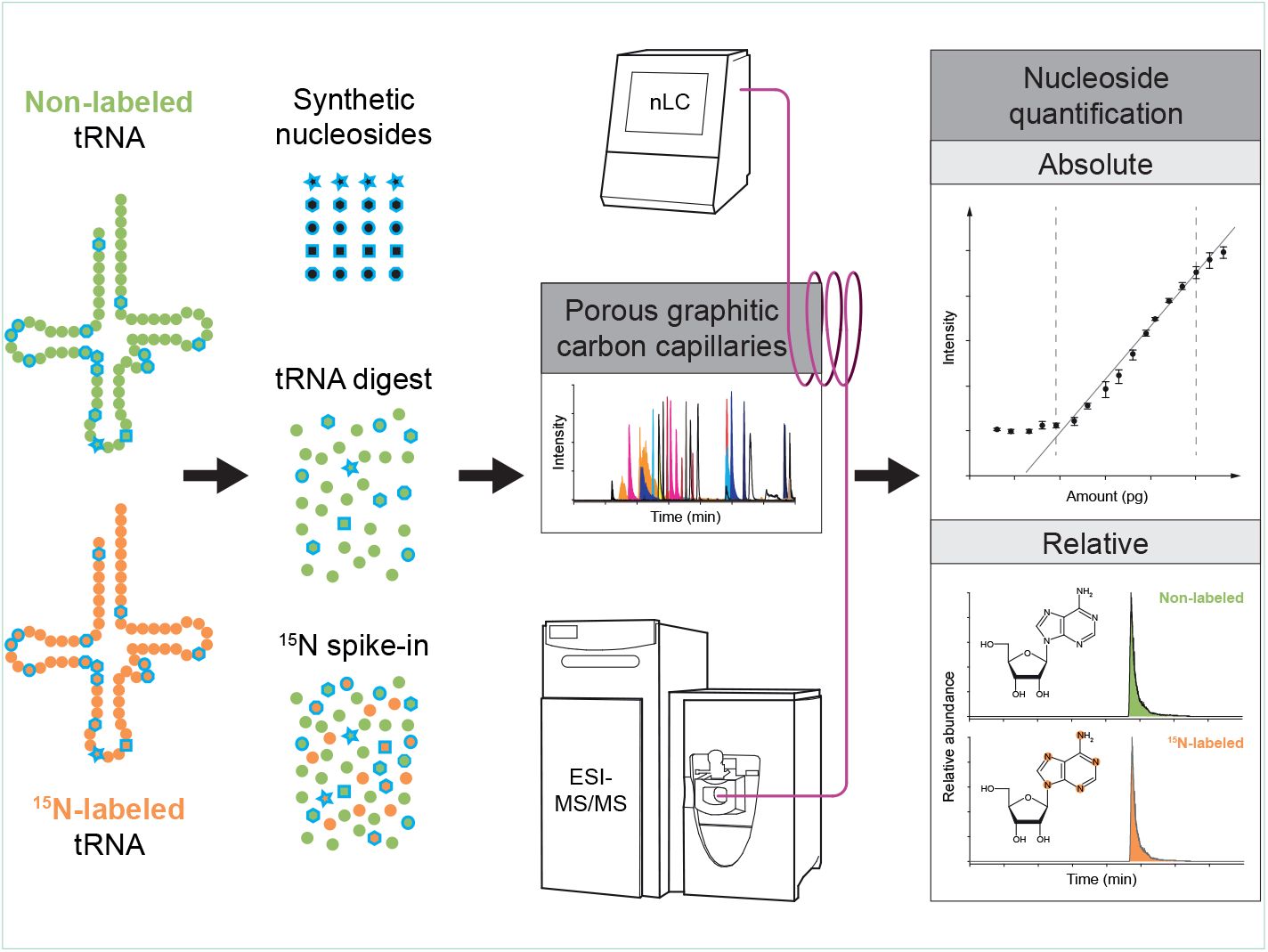

## REFERENCES

1. Shoemaker, C.J. and Green, R. (2012) Translation drives mRNA quality control. Nat. Struct. Mol. Biol., 19, 594–601.

2. Helm, M. and Alfonzo, J.D. (2014) Posttranscriptional RNA Modifications: playing metabolic games in a cell’s chemical Legoland. Chem. Biol., 21, 174–185.

3. Grosjean, H. (2009) Nucleic Acids Are Not Boring Long Polymers of Only Four Types of Nucleotides: A Guided Tour. In Grosjean, H. (ed.), DNA and RNA Modification Enzymes: Structure, Mechanism, Function and Evolution. Landes Bioscience, pp. 1–18.

4. Jackman, J.E. and Alfonzo, J.D. (2013) Transfer RNA modifications: nature’s combinatorial chemistry playground. Wiley Interdiscip. Rev. RNA, 4, 35–48.

5. Chan, C.T., Dyavaiah, M., DeMott, M.S., Taghizadeh, K., Dedon, P.C. and Begley, T.J. (2010) A quantitative systems approach reveals dynamic control of tRNA modifications during cellular stress. PLoS Genet., 6, e1001247.

6. Alings, F., Sarin, L.P., Fufezan, C., Drexler, H.C. and Leidel, S.A. (2015) An evolutionary approach uncovers a diverse response of tRNA 2-thiolation to elevated temperatures in yeast. RNA, 21, 202–212.

7. Nedialkova, D.D. and Leidel, S.A. (2015) Optimization of Codon Translation Rates via tRNA Modifications Maintains Proteome Integrity. Cell, 161, 1606–1618.

8. Grosjean, H., Breton, M., Sirand-Pugnet, P., Tardy, F., Thiaucourt, F., Citti, C., Barre, A., Yoshizawa, S., Fourmy, D., de Crecy-Lagard, V. et al. (2014) Predicting the minimal translation apparatus: lessons from the reductive evolution of mollicutes. PLoS Genet., 10, e1004363.

9. Laguesse, S., Creppe, C., Nedialkova, D.D., Prevot, P.P., Borgs, L., Huysseune, S., Franco, B., Duysens, G., Krusy, N., Lee, G. et al. (2015) A Dynamic Unfolded Protein Response Contributes to the Control of Cortical Neurogenesis. Dev. Cell, 35, 553–567.

10. Zinshteyn, B. and Gilbert, W.V. (2013) Loss of a conserved tRNA anticodon modification perturbs cellular signaling. PLoS Genet., 9, e1003675.

11. Sarin, L.P. and Leidel, S.A. (2014) Modify or die?--RNA modification defects in metazoans. RNA Biol., 11, 1555–1567.

12. Torres, A.G., Batlle, E. and Ribas de Pouplana, L. (2014) Role of tRNA modifications in human diseases. Trends Mol. Med., 20, 306–314.

13. Delaunay, S., Rapino, F., Tharun, L., Zhou, Z., Heukamp, L., Termathe, M., Shostak, K., Klevernic, I., Florin, A., Desmecht, H. et al. (2016) Elp3 links tRNA modification to IRES-dependent translation of LEF1 to sustain metastasis in breast cancer. J. Exp. Med., 213, 2503–2523.

14. Wei, F.Y., Suzuki, T., Watanabe, S., Kimura, S., Kaitsuka, T., Fujimura, A., Matsui, H., Atta, M., Michiue, H., Fontecave, M. et al. (2011) Deficit of tRNA(Lys) modification by Cdkal1 causes the development of type 2 diabetes in mice. J. Clin. Invest., 121, 3598–3608.

15. Chen, C., Tuck, S. and Byström, A.S. (2009) Defects in tRNA modification associated with neurological and developmental dysfunctions in *Caenorhabditis elegans* elongator mutants. PLoS Genet., 5, e1000561.

16. Suzuki, T., Nagao, A. and Suzuki, T. (2011) Human mitochondrial diseases caused by lack of taurine modification in mitochondrial tRNAs. Wiley Interdiscip. Rev. RNA, 2, 376–386.

17. Thuring, K., Schmid, K., Keller, P. and Helm, M. (2016) Analysis of RNA modifications by liquid chromatography-tandem mass spectrometry. Methods, 107, 48–56.

18. Ross, R., Cao, X., Yu, N. and Limbach, P.A. (2016) Sequence mapping of transfer RNA chemical modifications by liquid chromatography tandem mass spectrometry. Methods, 107, 73–78.

19. Russell, S.P. and Limbach, P.A. (2013) Evaluating the reproducibility of quantifying modified nucleosides from ribonucleic acids by LC-UV-MS. J. Chromatogr. B Analyt. Technol. Biomed. Life Sci., 923–924, 74–82.

20. Gehrke, C.W. and Kuo, K.C. (1989) Ribonucleoside analysis by reversed-phase high-performance liquid chromatography. J. Chromatogr., 471, 3–36.

21. Björk, G.R., Huang, B., Persson, O.P. and Byström, A.S. (2007) A conserved modified wobble 52 nucleoside (mcm^5^s^2^U) in lysyl-tRNA is required for viability in yeast. RNA, 13, 1245–1255.

22. Osipov, A.S. and Orlov, E.N. (2012) Separation of positional isomers using chiral chromatography columns. Pharm. Chem. J+, 46, 288–291.

23. Pereira, L. (2008) Porous Graphitic Carbon as a Stationary Phase in HPLC: Theory and Applications. J. Liq. Chromatogr. Relat. Technol., 31, 1687–1731.

24. Michel, M. and Buszewski, B. (2009) Porous graphitic carbon sorbents in biomedical and environmental applications. Adsorption, 15, 193–202.

25. Gundersen, J.L. (2001) Separation of isomers of nonylphenol and select nonylphenol polyethoxylates by high-performance liquid chromatography on a graphitic carbon column. J. Chromatogr. A, 914, 161–166.

26. De Matteis, C.I., Simpson, D.A., Euerby, M.R., Shaw, P.N. and Barrett, D.A. (2012) Chromatographic retention behaviour of monosubstituted benzene derivatives on porous graphitic carbon and octadecyl-bonded silica studied using molecular modelling and quantitative structure-retention relationships. J. Chromatogr. A, 1229, 95–106.

27. West, C., Elfakir, C. and Lafosse, M. (2010) Porous graphitic carbon: a versatile stationary phase for liquid chromatography. J. Chromatogr. A, 1217, 3201–3216.

28. Reepmeyer, J.C., Brower, J.F. and Ye, H. (2005) Separation and detection of the isomeric equine conjugated estrogens, equilin sulfate and Σ^8,9^-dehydroestrone sulfate, by liquid chromatography--electrospray-mass spectrometry using carbon-coated zirconia and porous graphitic carbon stationary phases. J. Chromatogr. A, 1083, 42–51.

29. Liu, Q., Shi, J., Zeng, L., Wang, T., Cai, Y. and Jiang, G. (2011) Evaluation of graphene as an advantageous adsorbent for solid-phase extraction with chlorophenols as model analytes. J. Chromatogr. A, 1218, 197–204.

30. He, X. and Kozak, M. (2012) Development of a liquid chromatography–tandem mass spectrometry method for plasma-free metanephrines with ion-pairing turbulent flow online extraction. Anal. Bioanal. Chem., 402, 3003–3010.

31. Chaimbault, P., Petritis, K., Elfakir, C. and Dreux, M. (2000) Ion-pair chromatography on a porous graphitic carbon stationary phase for the analysis of twenty underivatized protein amino acids. J. Chromatogr. A, 870, 245–254.

32. Takeuchi, T., Kojima, T. and Miwa, T. (2000) Ion Chromatography of Inorganic Anions on Graphitic Carbon as the Stationary Phase. J. High Resolut. Chromatogr., 23, 590–594.

33. Guenu, S. and Hennion, M.C. (1994) On-line sample handling of water-soluble organic pollutants in aqueous samples using porous graphitic carbon. J. Chromatogr. A, 665, 243–251.

34. Harris, E.H. (2009) Resources for the investigator. In Witman, E.H., Harris. D.B., Stern, G.B. (eds.), The Chlamydomonas Sourcebook (Second Edition). Academic Press, London, pp. 303–308.

35. Barth, J., Bergner, S.V., Jaeger, D., Niehues, A., Schulze, S., Scholz, M. and Fufezan, C. (2014) The interplay of light and oxygen in the reactive oxygen stress response of *Chlamydomonas reinhardtii* dissected by quantitative mass spectrometry. Mol. Cell. Proteomics, 13, 969–989.

36. Lecanda, A., Nilges, B.S., Sharma, P., Nedialkova, D.D., Schwarz, J., Vaquerizas, J.M. and Leidel, S.A. (2016) Dual randomization of oligonucleotides to reduce the bias in ribosome-profiling libraries. Methods, 107, 89–97.

37. Meiring, H.D., van der Heeft, E., ten Hove, G.J. and de Jong, A.P.J.M. (2002) Nanoscale LC–MS(n): technical design and applications to peptide and protein analysis. J. Sep. Sci., 25, 557–568.

38. Boccaletto, P., Machnicka, M.A., Purta, E., Piątkowski, P., Bagiński, B., Wirecki, T.K., de Crécy-Lagard, V., Ross, R., Limbach, P.A., Kotter, A. et al. (2017) MODOMICS: a database of RNA modification pathways. 2017 update. Nucleic Acids Res., doi:10.1093/nar/gkx1030.

39. Bald, T., Barth, J., Niehues, A., Specht, M., Hippler, M. and Fufezan, C. (2012) pymzML--Python module for high-throughput bioinformatics on mass spectrometry data. Bioinformatics, 28, 1052–1053.

40. Leufken, J., Niehues, A., Sarin, L.P., Wessel, F., Hippler, M., Leidel, S.A. and Fufezan, C. (2017) pyQms enables universal and accurate quantification of mass spectrometry data. Mol. Cell. Proteomics, 16, 1736–1745.

41. Martens, L., Chambers, M., Sturm, M., Kessner, D., Levander, F., Shofstahl, J., Tang, W.H., Rompp, A., Neumann, S., Pizarro, A.D. et al. (2011) mzML--a community standard for mass spectrometry data. Mol. Cell. Proteomics, 10, R110 000133.

42. Kessner, D., Chambers, M., Burke, R., Agus, D. and Mallick, P. (2008) ProteoWizard: open source software for rapid proteomics tools development. Bioinformatics, 24, 2534–2536.

43. Gower, J.C. (1971) A general coefficient of similarity and some of its properties. Biometrics, 27, 857–871.

44. Wan, Q.H., Shaw, N., Davies, M.C. and Barrett, D.A. (1995) Chromatographic behaviour of positional isomers on porous graphitic carbon. J. Chromatogr. A, 697, 219–227.

45. Agrofoglio, L.A., Bezy, V., Chaimbault, P., Delepee, R., Rhourri, B. and Morin, P. (2007) Mass spectrometry based methods for analysis of nucleosides as antiviral drugs and potential tumor biomarkers. Nucleosides Nucleotides Nucleic Acids, 26, 1523–1527.

46. Pabst, M., Grass, J., Fischl, R., Leonard, R., Jin, C., Hinterkorner, G., Borth, N. and Altmann, F. (2010) Nucleotide and nucleotide sugar analysis by liquid chromatography-electrospray ionization-mass spectrometry on surface-conditioned porous graphitic carbon. Anal. Chem., 82, 9782–9788.

47. Giessing, A.M., Scott, L.G. and Kirpekar, F. (2011) A nano-chip-LC/MSn based strategy for characterization of modified nucleosides using reduced porous graphitic carbon as a stationary phase. J. Am. Soc. Mass Spectrom., 22, 1242–1251.

48. Lim, C.K. (1989) Electronic interaction chromatography on porous graphitic carbon. Separation of [^99m^Tc]pertechnetate and perrhenate anions. Biomed. Chromatogr., 3, 92–93.

49. Bruckl, T., Globisch, D., Wagner, M., Muller, M. and Carell, T. (2009) Parallel isotope-based quantification of modified tRNA nucleosides. Angew. Chem. Int. Ed. Engl., 48, 7932–7934.

50. Kellner, S., Neumann, J., Rosenkranz, D., Lebedeva, S., Ketting, R.F., Zischler, H., Schneider, D. and Helm, M. (2014) Profiling of RNA modifications by multiplexed stable isotope labelling. Chem. Commun. (Camb), 50, 3516–3518.

51. Kellner, S., Ochel, A., Thuring, K., Spenkuch, F., Neumann, J., Sharma, S., Entian, K.D., Schneider, D. and Helm, M. (2014) Absolute and relative quantification of RNA modifications via biosynthetic isotopomers. Nucleic Acids Res., 42, e142.

